# Cyp26b1 is required for proper airway epithelial differentiation during lung development

**DOI:** 10.1101/678581

**Authors:** Edward Daniel, Gabrielle I. Sutton, Yadanar Htike, Ondine Cleaver

**Author notes:** Corresponding author: Ondine Cleaver, Department of Molecular Biology, University of Texas Southwestern Medical Center, 5323 Harry Hines Blvd., NA8.300, Dallas, Texas 75390-9148, USA.

## Abstract

Proper organ development depends on coordinated communication between multiple cell types. Retinoic acid (RA) is an autocrine and paracrine signaling molecule critical for the development of most organs including the lung. Both RA excess and deficiency lead to drastic alterations in embryogenesis, often culminating in embryonic or neonatal lethality. Therefore, RA levels must be spatially and temporally titrated to ensure proper organogenesis. Despite extensive work detailing the effects of RA deficiency in early lung morphogenesis, little is known about how RA levels are modulated during late lung development. Here, we investigate the role of the RA catabolizing protein Cyp26b1 in lung development. Cyp26b1 is highly enriched in lung endothelial cells (ECs) throughout the course of development. We find that loss of Cyp26b1 impacts differentiation of the distal epithelium without appreciably affecting proximal airways, EC lineages, or stromal populations. Cyp26b1^−/−^ lungs exhibit an increase in cellular density, with an expansion of distal progenitors at the expense of alveolar type 1 (AT1) cells, which culminates in neonatal death. Exogenous administration of RA in late gestation was able to partially reproduce this defect in epithelial differentiation; however, transcriptional analyses of Cyp26b1^−/−^ lungs and RA-treated lungs reveal separate, but overlapping, transcriptional responses. These data suggest that the defects observed in Cyp26b1^−/−^ lungs are caused by both RA-dependent and RA-independent mechanisms. This work highlights critical cellular crosstalk during lung development involving a crucial role for Cyp26b1-expressing endothelium, and identifies a novel RA rheostat in lung development.

**HIGHLIGHTS:**

- Cyp26b1 is highly expressed in lung ECs throughout development
- Cyp26b1-null lungs fail to undergo proper differentiation of distal epithelium leading to an increase in progenitors and AT2 cells at the expense of AT1 cells
- Functional and transcriptional analyses suggest both RA-dependent and RA-independent mechanisms

## INTRODUCTION

Organogenesis requires highly orchestrated crosstalk between endothelial cells (ECs), epithelial cells, and stromal cells in order to form a mature and functional organ. These cell types communicate with one another using a multitude of distinct signaling pathways that must be activated at the right place and time to promote proper development (Kraus and Grapin-Botton, 2012; Rankin et al., 2018). Coordination is central to development, and aberrations in any one step of this multistep process can have catastrophic developmental consequences. Mechanisms underlying spatiotemporal control of these signals over the course of organogenesis remains poorly understood.

Lung development begins at E9.0 with specification of lung progenitors along the ventral side of the anterior foregut endoderm (Herriges and Morrisey, 2014; Shi et al., 2009; Warburton et al., 2010). These progenitors undergo initial bud formation, followed by highly stereotyped and hierarchical branching to form the lung airway tree (Metzger et al., 2008). During this process, the lung epithelium segregates into 2 separate groups: the distal airways that give rise to alveoli where gas exchange occurs, and to proximal airways that form the bronchi and bronchioles. By E16.5, the distal airways begin to differentiate into pre-alveolar structures called canaliculi and saccules, consisting of alveolar type 2 (AT2) surfactant-producing cells and alveolar type 1 (AT1) gas-exchanging cells. These structures mediate gas exchange in neonates and full alveolarization occurs postnatally (Herriges and Morrisey, 2014; Morrisey and Hogan, 2010). In humans, failure to form early saccules results in respiratory distress syndrome (RDS), with a mortality rate as high as 50% depending on the size of the infant at birth (Gallacher et al., 2016).

Retinoic acid (RA) is a critical signaling molecule that exhibits highly regulated spatiotemporal control during embryogenesis. RA directs cell fate by binding to the RAR and RXR family of nuclear receptors to direct changes in gene transcription. RA is derived from Vitamin A (retinol) consumed in our diet. Vitamin A is first dehydrogenated to form retinaldehdyde, then retinaldehyde is further dehydrogenated by Raldh1-3 to form the active forms of RA, with all-trans retinoic acid (atRA) being the most abundant and potent form. Once synthesized, RA can act as an autocrine signal or can diffuse to nearby cells as a paracrine signal, forming a local gradient (Duester, 2008). Opposing RA signaling is the Cyp26 family of P450 enzymes, consisting of Cyp26a1, Cyp26b1, and Cyp26c1. These enzymes metabolize RA into inactive forms. Cells expressing P450 enzymes can act as a local sink for RA, thereby reducing local RA signaling. The expression of these genes require precise control as genetic or pharmacologic manipulations resulting in RA excess or deficiency drastically affects nearly every developing organ (Duester, 2008; Rhinn and Dolle, 2012). Spatiotemporal regulation of RA signaling can therefore be achieved through regulated expression of Raldh enzymes in RA-synthesizing cells and Cyp26 enzymes in RA-degrading cells to form local, tightly controlled RA gradients.

Lung development and maturation require proper spatiotemporal regulation of RA signaling. Mice lacking Raldh2 or fed on a Vitamin A-deficient diet fail to initiate lung formation, resulting in lung agenesis (Wang et al., 2006; Wilson et al., 1953). In the initial lung bud, RA activates the Wnt cascade via Shh, and inhibits TGF-β signaling (Chen et al., 2010; Chen et al., 2007; Rankin et al., 2016; Rankin et al., 2018). Together, this leads to upregulation of Fgf10, a critical factor necessary for lung bud formation and branching (Desai et al., 2004; Park et al., 1998; Wang et al., 2006). Once lung branching is initiated, RA activity decreases until birth. allowing proper epithelial branching and distal airway differentiation. Culturing E12.5 lung explants with RA led to a reduction in branching due to decreased Fgf10 (Malpel et al., 2000). Likewise, forcing increased RA signaling in distal lung epithelial cells through a constitutively active RAR *in vivo* prevents distal saccule differentiation (Wongtrakool et al., 2003). Lastly, a third, separate role for RA occurs postnatally during alveolar maturation. At this stage, RA promotes increased alveolar septation, alveolar number, and alveolar surface area (Massaro and Massaro, 1996, 2000; Massaro et al., 2003; McGowan et al., 2000). Taken together, these studies indicate that RA regulates several distinct steps during lung development in a highly temporally-defined manner.

Although RA signaling has been shown to regulate specification and differentiation and lung cell types, the role of RA-catabolizing Cyp26 enzymes during lung development remains unknown. Cyp26a1 is expressed in the epithelium during early stages of lung branching, but its expression is absent by E16.5 when distal epithelial differentiation begins (Malpel et al., 2000). On the other hand, Cyp26b1 is expressed throughout the lung mesenchyme, but not the epithelium, at E18.5 (Abu-Abed et al., 2002). Deletion of Cyp26b1 leads to neonatal lethality, which was suggested to occur from pulmonary dysfunction, although this phenotype was not characterized further (Yashiro et al., 2004). These data suggest Cyp26b1 may be required in the lung during late gestation to reduce RA signaling and promote proper distal epithelial differentiation.

Here, we identify a critical role for Cyp26b1 during lung organogenesis. Cyp26b1 is highly enriched in lung ECs throughout development, and loss of Cyp26b1 results in a delay in the formation of distal airways in late gestation. Cyp26b1^−/−^ lungs exhibit increased cellular density and contain an expansion of an early distal progenitor population at the expense of mature gas-exchanging AT1 cells. Exogenous administration of atRA during late gestation phenocopies loss of Cyp26b1, suggesting that the phenotype is due at least in part to excess RA. Transcriptional analysis of atRA-treated and Cyp26b1^−/−^ lungs however reveal only partially overlapping responses, suggesting additional signaling pathways downstream of Cyp26b1 in lung. These findings identify Cyp26b1 as a novel endothelial rheostat for RA activity in the developing lung and reveal RA-independent functions during lung development.

## MATERIALS AND METHODS

### Mice and embryo handling

Experiments were performed in accordance with protocols approved by the UT Southwestern Medical Center IACUC. Cyp26b1 mutant alleles were generated using CRISPR/Cas9 as previously described (Yang et al., 2013). In brief, Cas9 mRNA and in vitro transcribed sgRNAs were injected directly into C57BL/6J oocytes. sgRNA sequences used are Table S1. Successfully generated alleles were crossed into a CD1 background up to 5 backcrosses. Images of whole embryos and organs were taken with an iPhone XS (Apple) and NeoLumar stereomicroscope (Zeiss) using a DP-70 camera (Olympus), respectively.

RA administration to pregnant dams was performed as described previously (Cadot et al., 2012; Okano et al., 2012b). atRA (Sigma) was reconstituted in corn oil at 50 mg/mL. Pregnant dams were gavaged either 100 mg/kg atRA or equivalent dose of corn oil at E15.5, E16.5, and E17.5. Immediately prior to administration, atRA was resuspended to a consistent viscosity to ensure proper dosage. Reconstituted atRA was stored in the dark at 4C in between dosings for the same experiment. Different experiments used freshly prepared atRA suspension prepared immediately before the first dose.

### Measurements of embryo and lung weights

E18.5 embryos were dissected out of the uterus, blotted dry, and weighed. Wet and dry weights for the lungs were determined as previously described (Murata et al., 2007). Briefly, lungs were dissected out of embryos, blotted dry, and weighed on a pre-weighed piece of aluminum foil to determine the wet weight. Lungs on the aluminum foil were then dried overnight at 55°C and weighed again to determine the dry weight. Wet and dry weights were standardized to embryonic weight to determine relative wet and dry weights.

### Histologic and immunofluorescent analysis on sections

E12.5 – E18.5 embryos, lungs, and kidneys were dissected and fixed in 4% PFA/PBS overnight at 4°C. Tissues were washed in PBS the next day and dehydrated in a series of ethanol washes to 100% ethanol. Tissues were washed twice in 100% ethanol for 30 min (min), followed by 2 washes in xylene for 10 min each. Tissues were then moved to paraplast and washed at least 3 times in 100% paraplast at 60°C before incubating overnight at 60°C. Tissues were then mounted and sectioned at 10 μm on a microtome. For hematoxylin and eosin stains, sections were washed in xylene, 100% ethanol, and 95% ethanol 2 times for 3 min each. Slides were then kept under running water for 2 min followed by a 45 second incubation with hematoxylin (Gill’s Method, Fisher Chemical) and an additional 5 min under running water. Slides were submerged in acid alcohol (99 mL 70% ethanol + 1 mL 12N HCl) 4-5 times and washed under running water for 5 min. Slides were then submerged in 0.1% sodium bicarbonate for 1 min and washed for 5 min under running water. Eosin staining was performed by incubating slides with Eosin Y (Acros Organics) for 2 min followed by 3-4 1 min washes under running water. Lastly, slides were dehydrated to 100% ethanol, washed in xylene, and mounted using Permount. Images were taken using a Zeiss Axiovert 200M scope and a DP-70 camera (Olympus).

For immunofluorescent stains, slides were baked at 60°C for 10 min and allowed to cool down to room temperature before deparaffinizing in xylene 2 times for 5 min each and rehydrating through an ethanol series ending on brief wash in water. Once in aqueous solution, the stain was performed as described previously (Daniel et al., 2018). Briefly, slides were washed in PBS + 0.1% Triton-X 100 (Fisher Scientific, BP151-100) for 3 5-min washes. Slides were then treated with heat-mediated antigen retrieval in 1 μM Tris pH 7.5, 5 μM EDTA pH 8.0 prior to blocking depending on the antibody. Primary antibody incubations were done at 4°C overnight (for antibody information and dilutions, see Table S2). Slides were then washed in PBS, incubated in secondary antibody for 1 hr at room temperature. To reduce autofluorescence from red blood cells, slides were incubated in a 50 mM NH_4_Cl, 10 mM CuSO_4_, pH = 5.0 solution for 15 min followed by 2 5- min PBS + 0.1% Triton X-100 washes. Slides were then incubated with DAPI for 10 min, washed in PBS + 0.1% Triton X-100, and mounted using Prolong Gold Mounting Medium. Images were obtained using an A1R Nikon confocal microscope.

### Digoxigenin-labeled RNA probes and *in situ* hybridizations

Cyp26b1 full coding sequence clone was acquired from Dharmacon (BC059246) and linearized using EcoRI. Probe synthesis was performed as described previously (Daniel et al., 2018). Briefly, probes were synthesized at 37°C for 2-4 hrs in digoxigenin-synthesis reaction mixture with T3 RNA polymerase (Roche). After synthesis, DNA was eliminated by adding RQ1 DNase I (Promega) and RNA probes were purified using Micro Bio-spin columns (Bio-RAD). 10x hybridization stock was prepared at 10 μg/mL by adding the appropriate volume of pre-hybridization buffer.

*In situ* hybridizations were performed as described previously (Daniel et al., 2018). Briefly, E12.5 – E18.5 embryos, lungs and kidneys were fixed and embedded in paraffin as described above. Paraffin sections were de-paraffinized in xylene, then rehydrated to PBS before being treated with 15 μg/mL proteinase K for 15 min and fixed in 4% PFA. Slides were then washed and incubated with pre-hybridization buffer for 1 hr at room temperature before being hybridized with the specific probe at 1 μg/mL overnight at 65°C. Next day, slides were washed in 0.2x SSC then transferred to MBST before blocking with 2% blocking solution (Roche) for at least 1 hr at RT. Slides were then incubated with Anti-Dig alkaline phosphatase conjugated antibody (Roche, 1:4000) overnight at 4C. Next day, slides were washed in MBST 3x and NTMT 3x before incubating with BM purple (Roche) for color reaction. After color reaction, slides were fixed with 4% PFA and mounted using Permount. Images were taken using a Zeiss Axiovert 200M scope and a DP-70 camera (Olympus).

For fluorescent *in situ* hybridizations, the protocol is the same as the chromogenic assay through the 0.2x SSC wash on the second day. After that wash, slides were transferred to TNT and treated with 0.3% H_2_O_2_ for 30 min before washing in TNT 3x for 5 min. Slides were then blocked in 1% blocking reagent (Perkin Elmer) for at least 1 hour at room temperature. Slides were then incubated with Anti-Dig-Peroxidase (Roche, 1:500), rat anti-Endomucin (1:200), and rat anti-PECAM (1:200) overnight at 4C. After primary antibody incubation, slides were washed in TNT and treated with Fluorescein Amplification Reagent (Perkin Elmer, 1:50 in Plus Amplification Diluent) for 15 min at room temperature. Slides were then washed with TNT and incubated with donkey anti-rat 555 for 1 hour at room temperature. Lastly, slides were incubated with DAPI, washed in TNT and mounted using Prolong Gold Mounting Medium. Images were obtained using an A1R Nikon confocal microscope.

### RNA isolation and qRT-PCR

E18.5 lungs and kidneys were dissected and placed in RNAse-free Eppendorf tubes (Ambion) where they were manually dissociated using disposable plastic pestles. RNA extraction was performed using the RNeasy Mini Kit (Qiagen) following manufacturer’s instructions. mRNA concentrations were quantified on a NanoDrop 2000c Spectrophotometer (Thermo Fisher). RNA was standardized to the sample with the lowest concentration and was reverse transcribed with SuperScript III (Invitrogen) kits following manufacturer’s instructions using oligo-dTs. The resulting cDNA was diluted 1:6 in H_2_O prior to qRT-PCR analyses.

Transcripts were quantified using Power SYBR Green PCR Master Mix (Applied Biosciences) on a QuantStudio 3 Real-Time PCR System (Applied Biosciences). Primers used for qRT-PCR are listed in Table S1. Relative levels of transcripts were determining using the ΔΔCt method by first standardizing mean Ct for a given gene to the housekeeping gene Cyclophilin in the same sample and then calculating changes between samples. Data and statistical analyses were plotted and performed in GraphPad Prism 8.

### Flow cytometry

Primary lung cells were isolated from E16.5 – E18.5 lungs as previously described (Kim et al., 2005) using pan CD45-FITC, CD31-APC, Sca1 (Ly-6A)-APC-Cy7 (BD Pharmingen), EpCAM-PECy7 (BioLegend) with DAPI (Sigma) staining to eliminate dead cells. Briefly, whole E16.5 – E18.5 lungs were manually dissociated with a pestle in separate microcentrifuge tubes and then incubated with 2.5 mg/mL Collagenase A (Roche) and 20 ug/mL DNAse 1 (Sigma) for 45 min at 37°C on a nutator. Reaction was stopped by adding Wash media (PBS + 0.5% BSA + 2 mM EDTA + 1 mM CaCl_2_) to each reaction. Cells were pelleted and incubated with ACK Lysis buffer for 10 min on ice to lyse red blood cells. Next, cells were washed and filtered through 40 μm cell filters before incubating with the antibodies listed above. Cells were then washed again and resuspended with wash media + DAPI for analysis. Samples were analyzed on LSR II (BD Biosciences).

### Western blot

Protein extraction and western blot analyses were performed as previously described (Azizoglu et al., 2017). Briefly, E18.5 lungs were mechanically dissociated with a pestle and homogenized in PBS with 10 μg/mL aprotinin, 10 μg/mL leupeptin, and 10 μg/mL pepstatin. Triton X-100 was added to each tube to a final concentration of 1%. Samples were frozen at −80C overnight, thawed, and centrifuged at 10,000g for 5 min. Protein samples were quantified using Pierce BSA protein assay (ThermoScientific), standardized in Laemmli’s SDS-Sample Buffer (Boston Bioproducts). 40 μg of total protein from each lung lysate were run on a Western Blot. Antibodies and concentrations used are listed in Table S2.

### Quantification and Statistical Analysis

Cell counting for DAPI^+^, proSP-C^+^, Lamp3^+^, and Sox9^+^ cells was performed using Bitplane Imaris v.9.0.2. Cells were counted using the spots function to generate an initial count followed by manual editing to ensure proper counts. Lamp3^+^/proSP-C^+^ cells or Lamp3^+^/Sox9^+^ cells were quantified using the “colocalize spots” function in the Imaris XT analysis package. Images used in these quantifications encompass at least 5 random views of distal epithelium from at least 3 embryos per genotype and treatment condition. Images were captured to avoid all proximal epithelial structures that may improperly skew the results. Airspace area and number were calculated using FIJI. First, a threshold was applied to images of distal airways using the DAPI stain such that the areas occupied by cells were mostly filled. Next, the number and sizes of all blank spaces greater than 100 μm^2^ were calculated using the Analyze Particles function. The 100 μm^2^ threshold was chosen to eliminate false positive blank spaces that are due to gaps in between adjacent DAPI^+^ cells and not from actual open airspaces.

Data and statistical analyses were plotted and performed in GraphPad Prism 8 for all experiments described. Significance for cell quantifications, qRT-PCR analysis, and flow cytometry was determined using two-way ANOVA with Sidak multiple comparison test. Significance for embryo and lung weights was determined using one-way ANOVA with Tukey multiple comparison test.

## RESULTS

### Cyp26b1 is highly enriched in lung and kidney ECs

We previously identified *Cyp26b1* as an endothelial-enriched gene in embryonic organs, including kidney and lung, using RNA-sequencing (Daniel et al., 2018). Cyp26b1 was expressed in the mesenchyme of these embryonic organs, suggesting possible lung EC enrichment (Abu-Abed et al., 2002). To differentiate between the two major cell types resident in the lung mesenchyme - lung ECs and stromal cells - we assessed Cyp26b1 expression in publically available single cell RNA-sequencing (scRNA-seq) datasets of fetal, early postnatal, and adult tissues. These studies confirmed increased expression of Cyp26b1 in ECs compared to non-ECs (**Supplemental Fig. 1**) (Du et al., 2015; Du et al., 2017; Guo et al., 2019; Hochane et al., 2019; Lindstrom et al., 2018; Sabbagh et al., 2018; Tabula Muris et al., 2018).

To validate these data, we performed in situ hybridization for Cyp26b1 in E12.5, E15.5, and E18.5 embryonic organs. These experiments revealed an endothelial-like expression pattern in both lung and kidney (**Fig. 1A, E**). Consistent with previously published data, Cyp26b1 was also expressed in the developing limbs, the face and palate, the tongue, the hindbrain, intersomitic regions along the back, in endocardial cushions extending into the great vessels, and the epicardium (**Supplemental Fig. 2**) (Abu-Abed et al., 2002; Spoorendonk et al., 2008). By E15.5, Cyp26b1 expression is lungs was restricted to the ECs in the distal periphery (**Fig. 1A-D**). Likewise, Cyp26b1 expression in the kidney became restricted to ECs in the outer cortex, which surrounds developing nephrons (**Fig. 1E-H**). We validated EC specificity of Cyp26b1 using fluorescent in situ hybridization (FISH) analysis and co-staining with the endothelial markers PECAM and Endomucin (Emcn) (**Fig. 1D, H**). Of note, Cyp26b1^+^ punctae were observed in some epithelial and stromal regions, but at much lower levels compared to ECs (white arrows). Thus, Cyp26b1 expression is highly enriched in lung and kidney ECs throughout development.

**Figure 1.**
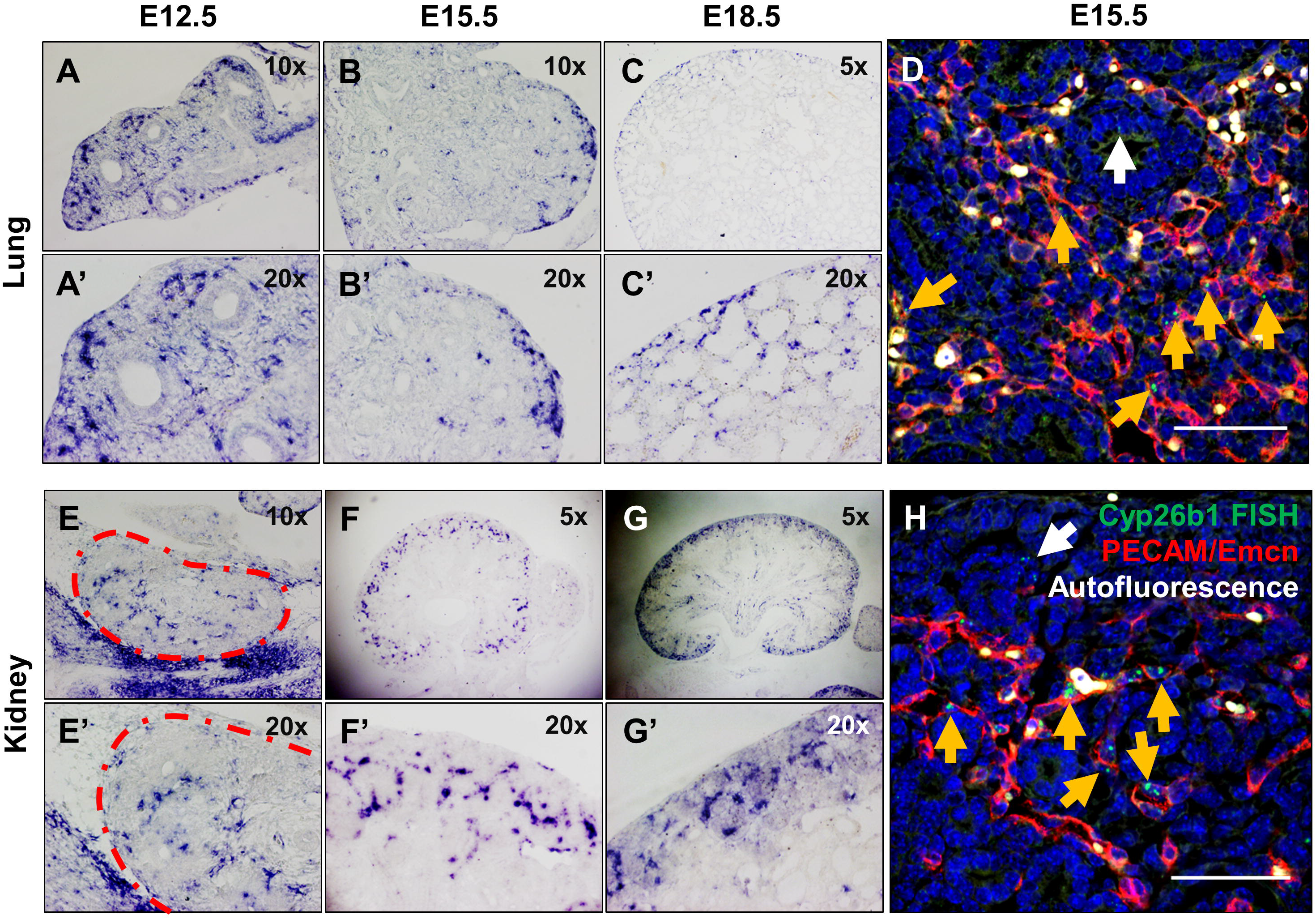
Cyp26b1 is highly enriched in lung and kidney endothelial cells during development. In situ hybridization of Cyp26b1 in the kidney (A-D) and lung (E-H) at E12.5 (A, E), E15.5 (B, D, F, H), and E18.5 (C, G). Magnifications for chromogenic assays shown. D,H) Fluorescent in situ hybridization for Cyp26b1 (green) co-stained with PECAM and Emcn (red) to mark ECs. Orange arrows are Cyp26b1^+^ punctae in PECAM/Emcn^+^ ECs. White arrow identifies Cyp26b1^+^ puncta in non-EC cell. Scale bar = 50 μm.

### Generation of Cyp26b1 null mice using CRISPR/Cas9

Previous work using a Cyp26b1 germline deficient mouse model showed that Cyp26b1-null mutants die shortly after birth due to respiratory distress (Yashiro et al., 2004). We generated two independently derived Cyp26b1-null mouse models using CRISPR/Cas9 to further explore this defects. Our first model contains an in-frame 2.6kb deletion from Exon 3 to Exon 6 (referred to as Cyp26b1^−^) and the second contains a 10bp deletion in Exon 3 leading to a frame-shift mutation (referred to as Cyp26b1^Δ10^). Both Cyp26b1^−/−^ and Cyp26b1^Δ10/Δ10^ embryos exhibited many of the same developmental defects previously described, including limb defects, craniofacial abnormalities, micrognathia, cleft palate, skin abnormalities, and spleen hypoplasia, although edema and hemorrhages were not observed at this stage as previously described (**Fig. 2A-B**) (Bowles et al., 2014; Dranse et al., 2011; Lenti et al., 2016; Okano et al., 2012a; Okano et al., 2012b; Yashiro et al., 2004). Additionally, Cyp26b1^−/−^ mice died shortly after birth and exhibited signs of respiratory distress, including air hunger mirroring the previously generated null model. These observations and previously published data suggest that lung development in Cyp26b1-null mice is abrogated in late gestation.

**Figure 2.**
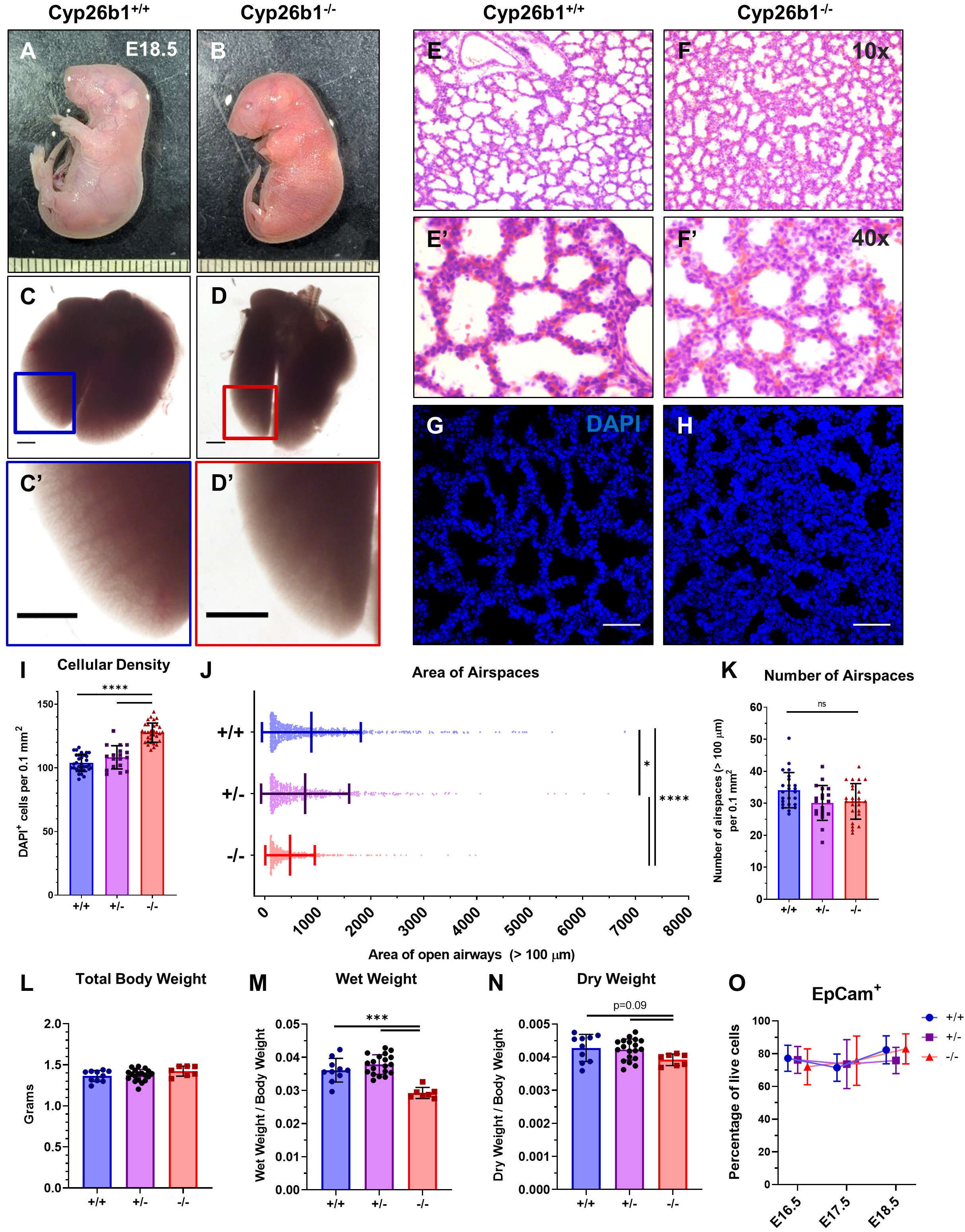
Cyp26b1 mutant lungs exhibit increased cellular density and decreased airspaces at late gestation. A-B) E18.5 Cyp26b1^+/+^ and Cyp26b1^−/−^ embryos at dissection. C-D) Lungs from Cyp26b1^+/+^ and Cyp26b1^−/−^ embryos. Zoomed areas in C’ and D’ highlight loss of distal airspaces in Cyp26b1^−/−^ lungs. E-F) H&E stain of E18.5 Cyp26b1^+/+^ and Cyp26b1^−/−^ lungs at 10x (E, F) and 40x (E’, F’) demonstrating increased septal wall thickness and smaller airspaces. G-H) Representative images of DAPI stains used in quantifications for I-K. I) Number of DAPI^+^ cells per 0.1 mm^2^ demonstrating increased cellularity in the mutants. J) Area of open airspaces (>100 μm^2^) in Cyp26b1^+/+^, Cyp26b1^+/−^, and Cyp26b1^−/−^ lungs. K) The total number of open airspaces (>100 μm^2^) per 0.1 mm^2^. L) Total body weight of E18.5 embryos. M) Weight of lungs immediately after dissection (wet) relative to total body weight in E18.5 embryos. N) Weight of lungs after drying relative to total body weight in E18.5 embryos. O) Flow cytometry analysis of EpCam^+^/CD31^−^/CD45^−^ cells in E16.5 – E18.5 lungs. Scale bars = 1 mm (A-D), 50 μm (G-H). Significance was determined using one-way ANOVA with Tukey multiple comparison test. \**P*<0.05, \*\*\**P*<0.001, \*\*\*\**P*<0.0001.

### Cyp26b1 null mice exhibit lung defects

To further explore how loss of Cyp26b1 affects lung development in late gestation, we analyzed Cyp26b1^+/+^ and Cyp26b1^−/−^ lungs at E18.5, immediately prior to birth. E18.5 Cyp26b1^−/−^ lungs were grossly smaller with decreased distal airspaces compared to control littermates but exhibited normal lobation (**Fig. 2C-D**). In tissue sections, Cyp26b1^−/−^ lungs appeared hypercellular, with increased septal wall thickness and smaller airspaces although the total number of airspaces was not different (**Fig. 2E-K**). To determine whether the phenotype is due to an increase in cell number or an increase in density of the same number of cells but failure of epithelial lumenogenesis, we measured the wet and dry weights of these lungs. Measurements of the wet and dry weight standardized to total body weight confirm that Cyp26b1^−/−^ lungs are both smaller at dissection without an increase in total cellular mass (**Fig. 2L-N**). To further assess changes in epithelial volume, we performed flow cytometry on E16.5 – E18.5 lungs with the epithelial marker EpCam. This analysis indicated that the number of epithelial cells did not differ between Cyp26b1^+/+^ and Cyp26b1^−/−^ lungs, at all 3 time points (**Fig. 2O**). These data suggest that the phenotype in Cyp26b1^−/−^ lungs is due to an increase in cellular density and not due to increased total cellular mass.

### Cyp26b1 is required for distal airway morphogenesis in late gestation

Our results show that distal airspaces were reduced in Cyp26b1^−/−^ lungs. To see when this phenotype arose, we analyzed earlier stages of lung development. Distal airway differentiation begins at E16.5 when the columnar epithelium undergoes dramatic cell shape changes that lead to lumen expansion and formation of distal saccules for gas exchange (Herriges and Morrisey, 2014; Shi et al., 2009; Warburton et al., 2010). Consistent with this timeline, Cyp26b1^−/−^ lungs begin to show gross morphological defects beginning at E16.5 characterized by increased cellular density and decreased distal airspace formation (**Supplemental Fig. 3**). During distal epithelial differentiation, epithelial cells undergo morphogenetic changes from a columnar, glandular-like structure to a flattened or rounded morphology for AT1 or AT2 cells, respectively (**Fig. 3A, C, E**). In Cyp26b1^−/−^ lungs, the epithelium exhibits a shift towards rounded epithelium while the transition to flattened epithelium appears delayed and epithelial cells remain relatively rounded (**Fig. 3B, D, F**). Although distal epithelial lumens in the mutant lungs are present at E16.5, these lumens are appreciably smaller and are only open in regions closest to the end of the proximal epithelium (**Fig. 3G, H**, magenta outlines).

**Figure 3.**
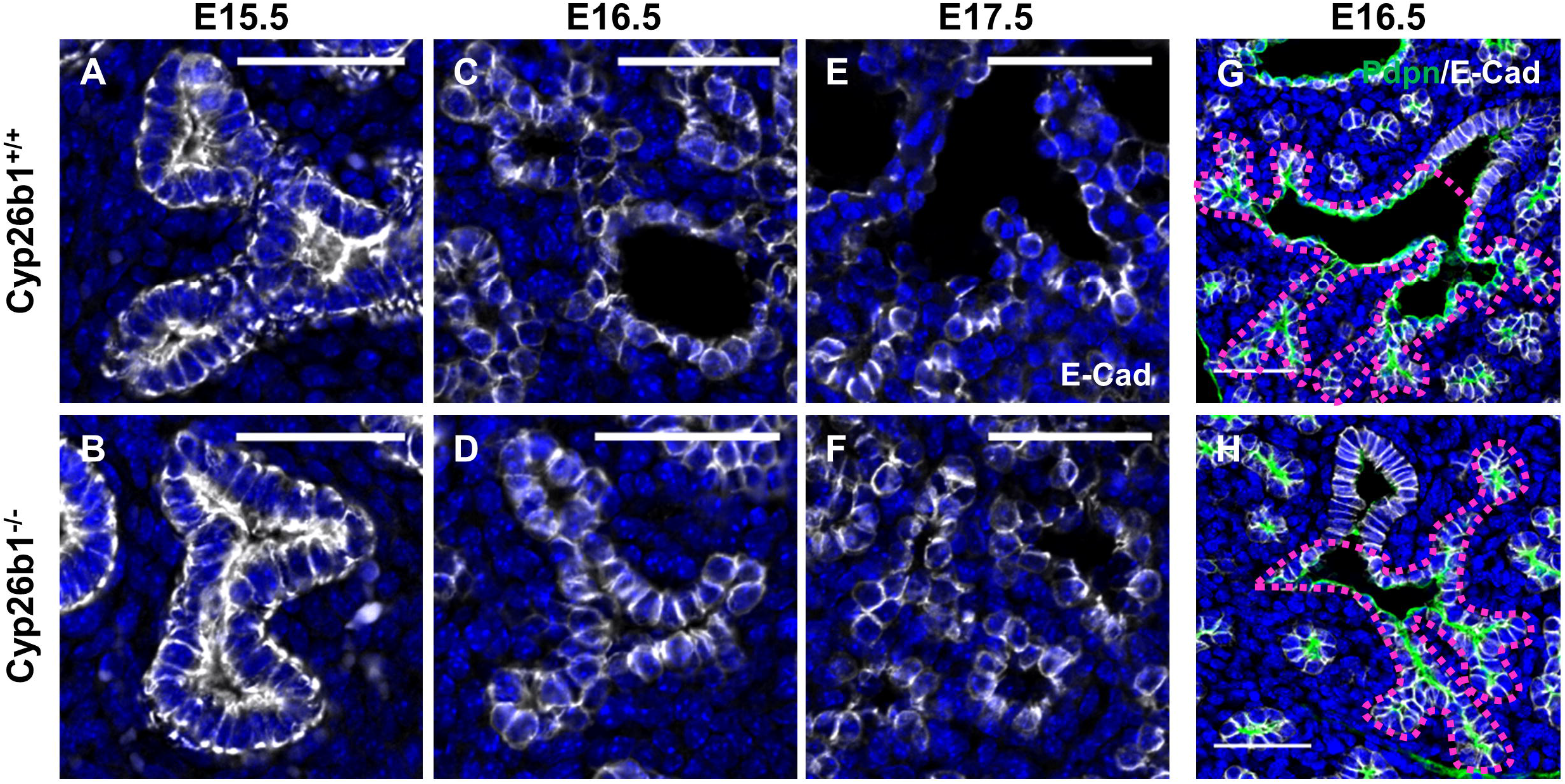
Defects in epithelial morphogenesis in Cyp26b1−/− lungs arise at E16.5. A-F) H&E of Cyp26b1^+/+^ and Cyp26b1^−/−^ lungs at E15.5 (A-B), E16.5 (C-D), and E17.5 (E-F). G-L) Immunofluorescent stains for E-Cad (white) to mark the epithelial structure at E15.5 (G,H), E16.5 (I,J), and E17.5 (K,L). M-N) Proximal to distal epithelial transition in E16.5 lungs stained with the AT1 cell marker Pdpn (green) and E-cad (white). Magenta outlines mark distal epithelial tree extending from the terminal bronchiole. Scale bar = 50 μm.

### Cyp26b1 is necessary for distal epithelial maturation

Based on these changes in epithelial morphology, we asked whether distal epithelial differentiation was affected in Cyp26b1^−/−^ lungs. Prior to E16.5, distal epithelial progenitor cells co-express Sox9, pro-Surfactant Protein C (proSP-C) and Aquaporin 5 (Aqp5) (Herriges and Morrisey, 2014; Shi et al., 2009; Warburton et al., 2010; Yang et al., 2016). At E16.5 coincident with morphogenetic changes, these distal epithelial progenitor cells differentiate into the two primary lineages: proSP-C ^+^ AT2 cells and Aqp5^+^ AT1 cells (**Fig.4A**).

Immunofluorescent stains for proSP-C at E18.5 reveal that Cyp26b1^−/−^ lungs contain an increase in AT2 cells compared to control littermates (**Fig. 4B-D**). Of note, Cyp26b1^−/−^ lungs contain a higher proportion of AT2 cells after standardizing for the increased number of DAPI^+^ cells per given area. Because proSP-C is not exclusively expressed in AT2 cells during development (Frank et al., 2019), we repeated the analysis utilizing the AT2-cell specific marker, Lamp3 (“Lysosomal associated membrane protein 3” or DC-Lamp). Quantification of AT2 cells using Lamp3 demonstrated a similar increase in Lamp3^+^ AT2 cells as compared to proSP-C^+^ AT2 cells in Cyp26b1^−/−^ lungs, with the vast majority of AT2 cells expressing both proteins (**Supplemental Fig. 4A-F**).

**Figure 4.**
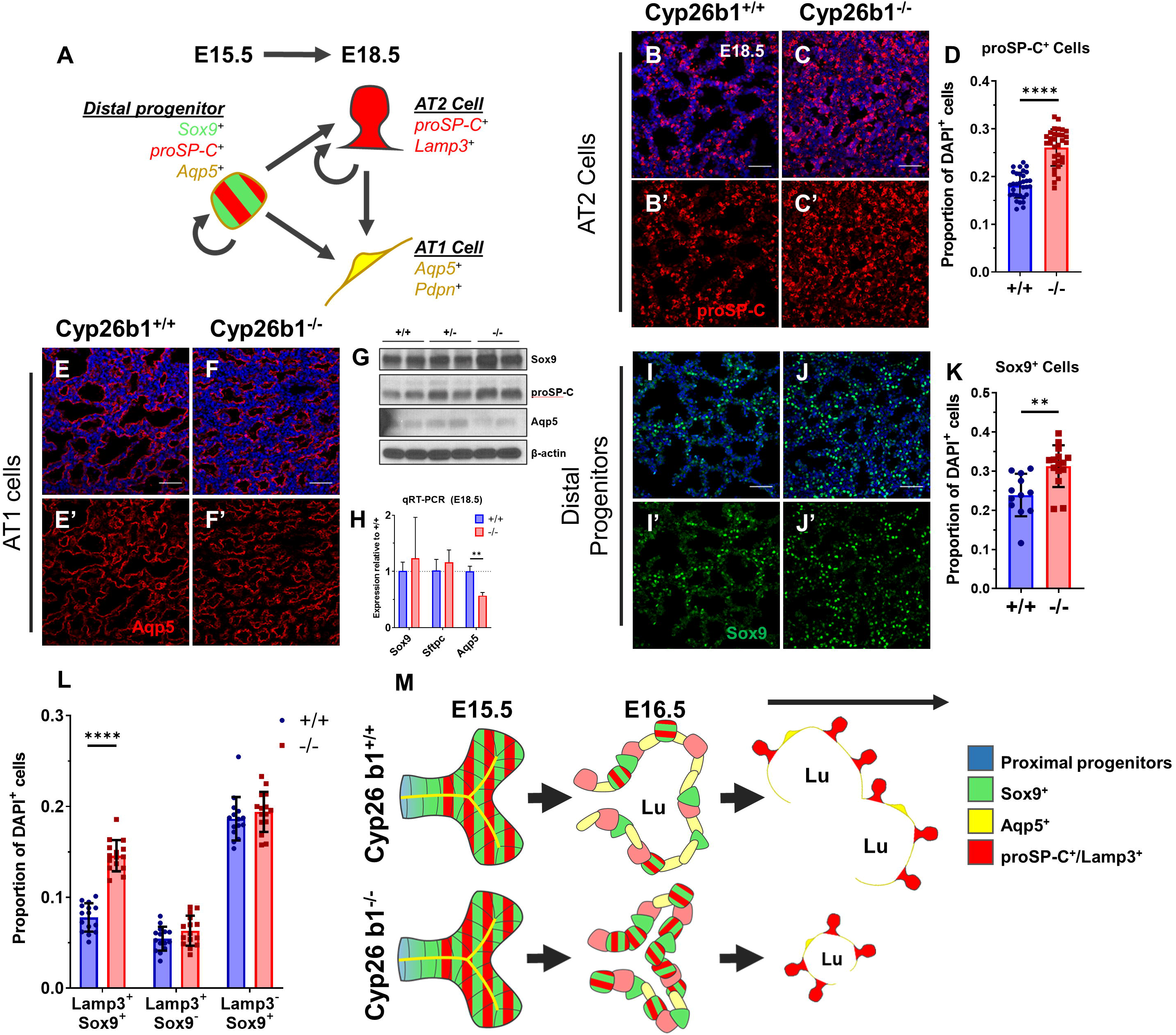
Distal epithelial progenitors and AT2 cells are expanded at the expense of AT1 cells in Cyp26b1^−/−^ lungs. A) Model of distal epithelial maturation. Distal progenitor cells express Sox9, proSP-C and Aqp5. These cells give rise to either AT1 or AT2 cells. B-D) IF stain for the AT2 marker proSP-C. (B-C) and quantification of cell counts (D). E, F) IF stain for Aqp5 to mark AT1 cells. G) Western blot analysis of Sox9, proSP-C, and Aqp5 in E18.5 lungs with β-actin as loading control. H) qRT-PCR of Sox9, Sftpc, and Aqp5 at E18.5. I-K) IF stain for the distal progenitor cell marker Sox9 (I, J) and quantification of cell counts (K). L) Quantification of Lamp3/Sox9 double positive cells (first set), Lamp3 single positive cells (second set), and Sox9 single positive cells (third set) as a proportion of DAPI^+^ cells. Note that the quantifications in D, K and L are standardized to the total number of DAPI^+^ cells. M) Model of distal epithelial maturation between Cyp26b1^+/+^ and Cyp26b1^−/−^ lungs. At E15.5 before Sox9^+^/proSP-C^+^/Aqp5^+^ distal progenitors (green, red, yellow respectively) begin to differentiate, the epithelium resembles a glandular-like structure. Beginning at E16.5, these tips will undergo morphologic changes from a columnar to a squamous epithelium and start forming early AT1 (yellow) and AT2 (red) cells. Concurrently, lumens (L) open at this stage forming the early saccules needed for gas exchange. These cells continue to mature for the rest of embryonic development and postnatally to form the mature functioning alveoli. In Cyp26b1^−/−^ lungs, epithelial development is not affected through E15.5. Once distal differentiation commences, the epithelium is able to alter its morphology towards a more squamous epithelium; however, lumens fail to fully open and the proportion of distal progenitors and AT1 cells have shifted. This ultimately results in fewer functional gas-exchanging units with smaller lumens, culminating in neonatal demise. Scale bar = 50 μm. Significance in D, K and L was determined using Student’s T-Test. Significance in H was determined using two-way ANOVA with Sidak multiple comparison test. \*\**P*<0.01, \*\*\*\**P*<0.0001.

We next assessed for changes in AT1 cells. Immunofluorescent stains for Aqp5 AT1 cells revealed that distal airway spaces are decreased in the mutants (**Fig. 4E, F**). Likewise, Podoplanin (Pdpn), an additional AT1 cell marker, mirrored the phenotypes seen with Aqp5 by immunofluorescence (**Supplemental Fig. 4G, H**). To estimate the relative abundance of AT1 cells, we performed western blot and qRT-PCR analyses on E18.5 tissues. Western blot analyses demonstrated a decrease in the AT1 cell marker Aqp5 and an increase in proSP-C, validating the immunofluorescent data (**Fig. 4G**). Likewise, qRT-PCR for the AT1 cell markers Aqp5, Pdpn, HOPX, and Ager/RAGE revealed ~40% decrease in mRNA abundance in Cyp26b1^−/−^ lungs; however, Sox9, Sftpc, and Lamp3 mRNA levels were not significantly altered (**Fig. 4H**, **Supplemental Fig. 4I**).

These apparent changes in epithelial differentiation led us to ask whether there were changes in the progenitor populations. Quantification of Sox9^+^ distal epithelial cells mirrored the results of proSP-C (**Fig. 4I-K**). To determine whether the increase in Sox9^+^ and proSP-C^+^/Lamp3^+^ cells in Cyp26b1^−/−^ lungs represent an increase in 2 distinct populations or in the same early distal progenitor population, we co-stained E18.5 lungs with both Sox9 and Lamp3. Quantification of these populations revealed an expansion in only Sox9^+^/Lamp3^+^ cells, with no change in Sox9^+^/Lamp3^−^ or Sox9^−^/Lamp3^+^ cells (**Fig. 4L** and **Supplemental Fig. 4J, K**). Therefore, Cyp26b1^−/−^ lungs contain an expanded early distal progenitor population.

### Defects in lung development are not due to off-target CRISPR lesions

Because our Cyp26b1 mutant mice were generated using CRISPR/Cas9-mediated mutagenesis, we considered the possibility that there may be off-target effects contributing to these phenotypes. These off-target effects can be eliminated by crossing two independently derived null alleles, to create compound heterozygotes. Any phenotypes observed in compound heterozygotes can then be fully attributed to loss of the target gene. We generated compound heterozygotes of Cyp26b1 by crossing the Cyp26b1^−^ allele with the Cyp26b1^Δ10^. E18.5 Cyp26b1^−/Δ10^ embryos completely phenocopied Cyp26b1^−/−^ embryos including gross developmental defects, decreased distal airspaces, increased cellularity, and increased relative numbers of distal epithelial progenitors and AT2 cells (**Supplemental Fig. 5**). Taken together, these data demonstrate that loss of Cyp26b1 result in changes in distal epithelial morphology and differentiation (**Fig. 4M**). More specifically, distal epithelial progenitor and AT2 cells are increased at the expense of AT1 cells in both models.

### Proximal airways, stroma, endothelia and lymphatics are unaffected in Cyp26b1^−/−^ lungs

We further assayed for changes in other populations in Cyp26b1^−/−^ lungs. Proximal airways in Cyp26b1^−/−^ lungs appeared unaffected as there was no appreciable difference in gross morphology or in the abundance of CCSP^+^ secretory cells (Clara cells) and Foxj1^+^ ciliated cells (bronchial in embryonic lungs) (**Supplemental Fig. 6A, B**). CCSP^+^, Sca1^+^ Bronchioalveolar Stem Cells (BASCs) are a stem cell population that reside at the transition between proximal and distal airways and can give rise to distal epithelium in the adult during lung regeneration (Kim et al., 2005; Lee et al., 2014). Analysis of BASCs by flow cytometry revealed no differences in this population at any stage in late gestation (**Supplemental Fig. 6C, D**).

Several groups have established a link between RA signaling and proper vascular formation in the heart through directing proper vascular smooth muscle cell differentiation and association of the vasculature with these smooth muscle cells (Braitsch et al., 2012; Wang et al., 2018; Xiao et al., 2018). We analyzed the vasculature and associated stroma in Cyp26b1^−/−^ lungs to see if similar changes can be identified. Gross analysis by IF of the vasculature and stroma revealed no overt changes (**Supplemental Fig. 7A-D**). Co-stains for VE-Cad and Pdgfr-β to mark the ECs and pericytes, respectively, did not reveal any differences in EC-pericyte coupling (**Supplemental Fig. 7E-F**). We also observed no differences in arterial, venous, and lymphatic development (**Supplemental Fig. 7G-J**). Lastly, RA has been shown to direct the differentiation of stromal-derived smooth muscle cells during lung organogenesis (Chen et al., 2014). Analysis using the marker Sm22a (protein product of *Tagln*) revealed that smooth muscle cells around both airways and vessels were not appreciably altered in Cyp26b1^−/−^ lungs (**Supplemental Fig. 7K-L**). These data show that formation of the vasculature, stroma, and associated lineages is not affected in Cyp26b1^−/−^ lungs.

### Cyp26b1 mutant lungs exhibit a partial RA response

Cyp26b1 dampens RA signaling by catabolizing RA into an inactive metabolite; therefore, we assessed levels of RA signaling in lungs lacking Cyp26b1. Because the main mechanism of RA is to modulate gene transcription, we assayed for genes involved in RA metabolism by qRT-PCR, several of which are direct targets of RA targets (RARβ, Stra6, Crabp2, Rbp1, Cyp26 family) (Balmer and Blomhoff, 2002; Rhinn and Dolle, 2012; Ross and Zolfaghari, 2011; Wu and Ross, 2010). Transcriptional analyses of these genes in whole lung lysates revealed upregulation of RARβ and Stra6, downregulation of the RA-synthesizing enzymes Raldh1 and Raldh3, and a trend towards increased and decreased expression of Crabp2 and Raldh2, respectively (**Fig. 5A**). Assaying the Cyp26 family, we first verified the mutation by assessing Cyp26b1 expression utilizing primers within the deleted region spanning the intron between Exon 4 and Exon 5 (**Fig. 5B**, “Cyp26b1-E4-5”). Repeating the PCR using primers outside of the deleted region demonstrated that Cyp26b1 is strongly upregulated in Cyp26b1^−/−^ lungs (**Fig. 5B**). Expression levels of Cyp26a1 and Cyp26c1 were unaffected (**Fig. 5B**). F/ISH for Cyp26b1 in mutant lungs validated the qRT-PCR data and revealed that the upregulation of Cyp26b1 came specifically from ECs (**Fig. 5C-H**). These data are consistent with our proposed model that RA signaling is increased in Cyp26b1^−/−^ lungs.

**Figure 5.**
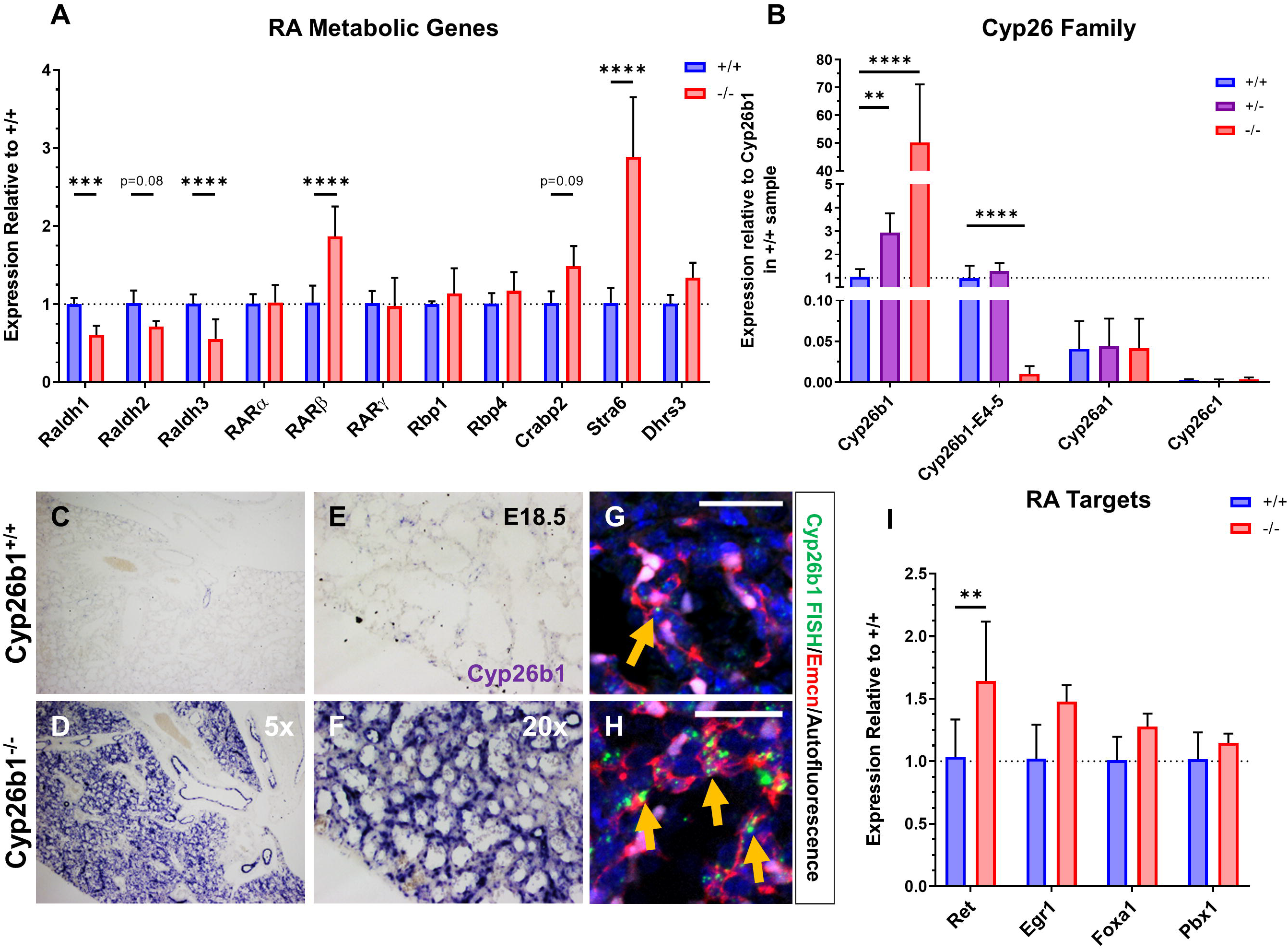
Cyp26b1^−/−^ lungs exhibit a mixed RA response. A) qRT-PCR for several members of RA metabolism in whole E18.5 lungs. B) qRT-PCR for the three Cyp26 family members all standardized to the expression of Cyp26b1 in Cyp26b1^+/+^ samples. Cyp26b1 utilizes primers outside of the deleted region (specifically spanning the intron between Exons 2 and 3) in Cyp26b1^−^ line whereas Cyp26b1-E4-5 utilizes primers designed within the deleted region (specifically spanning the intron between Exons 4 and 5). C-F) ISH for Cyp26b1 in E18.5 Cyp26b1^+/+^ (C,E) and Cyp26b1^−/−^ (D,F) lungs at 5x (C-D) and 20x (E-F) zoom. G-H) FISH for Cyp26b1 (green) costained with Emcn (red). Orange arrows mark Cyp26b1^+^ punctae in ECs. I) qRT-PCR for Ret, Egr1, Foxa1, and Pbx1 in whole E18.5 lungs. Scale bar = 25 μm (G-H). Significance was determined using two-way ANOVA with Sidak multiple comparison test. \*\**P*<0.01, \*\*\**P*<0.001, \*\*\*\**P*<0.0001.

We next asked whether other targets of RA signaling are altered in Cyp26b1^−/−^ lungs. We first analyzed expression levels of Ret, Egr1, Foxa1, and Pbx1, which have all been shown to be direct targets of RA signaling in other contexts (Balmer and Blomhoff, 2002; Probst et al., 2011; Rhinn and Dolle, 2012; Wong et al., 2012). qRT-PCR analyses of whole lung lysates for these genes revealed significant upregulation of Ret but not the other 3 genes (**Fig. 5I**). We asked whether other genes may be differentially regulated. During early lung morphogenesis, RA regulates multiple signaling pathways, including Fgfs, Tgf-β/Bmps, Wnts, and Shh (Chen et al., 2007; Desai et al., 2004; Malpel et al., 2000; Rankin et al., 2016; Rankin et al., 2018). Interestingly, qRT-PCR analysis for all of these pathways showed no change, except for downregulation of the Tgf-β target Tgfbi (**Supplemental Fig. 8**). Taken together, these data indicate that loss of Cyp26b1 can lead to a partial RA response in the lung in which genes involved in RA metabolism are transcriptionally altered but other established downstream targets are unaffected.

### RA partially contributes to the Cyp26b1^−/−^ phenotype in the lung

We sought to more clearly determine if RA is sufficient to induce the changes seen in the Cyp26b1^−/−^ lungs. Previous reports showed that exogenous administration of all trans retinoic acid (atRA) can partially phenocopy the defects in limb development, palate fusion, and skin barrier formation that are seen in Cyp26b1^−/−^ embryos (Cadot et al., 2012; Okano et al., 2012b). Following a similar dosing regimen, we gavaged 100 mg/kg atRA to Cyp26b1^+/−^ pregnant dams at E15.5, E16.5, and E17.5, dissected out the lungs at E18.5, and assessed for changes in lung development (**Fig. 6A**). 100% of Cyp26b1^−/−^ and ~30% of Cyp26b1^+/−^ and Cyp26b1^+/+^ embryos receiving exogenous atRA died in utero while the remainder died shortly after birth (**Fig. 6B**). Because all atRA-treated Cyp26b1^−/−^ embryos died by E18.5, we focused on the lungs of atRA-treated Cyp26b1^+/+^ and Cyp26b1^+/−^ embryos for these analyses. atRA-treated Cyp26b1^+/+^ and Cyp26b1^+/−^ lungs exhibited a loss of distal airspaces and increased cellular density that were indistinguishable from control Cyp26b1^−/−^ lungs (**Fig. 6C-L, W**, **Supplemental Fig. 9A-E**). Sox9^+^ distal progenitor cells were relatively more abundant in atRA-treated lungs per given area compared to their matched controls, but not to the same degree as that in Cyp26b1^−/−^ lungs (**Fig. 6M-Q, X**). Interestingly, proSP-C^+^ AT2 cells were only proportionally increased in atRA-treated Cyp26b1^+/−^ lungs, but not those of Cyp26b1^+/+^ (**Fig. 6R-V, Y**). These data show that excess RA is sufficient to induce morphologic changes in Cyp26b1^−/−^ lungs but does not fully recapitulate loss of Cyp26b1.

**Figure 6.**
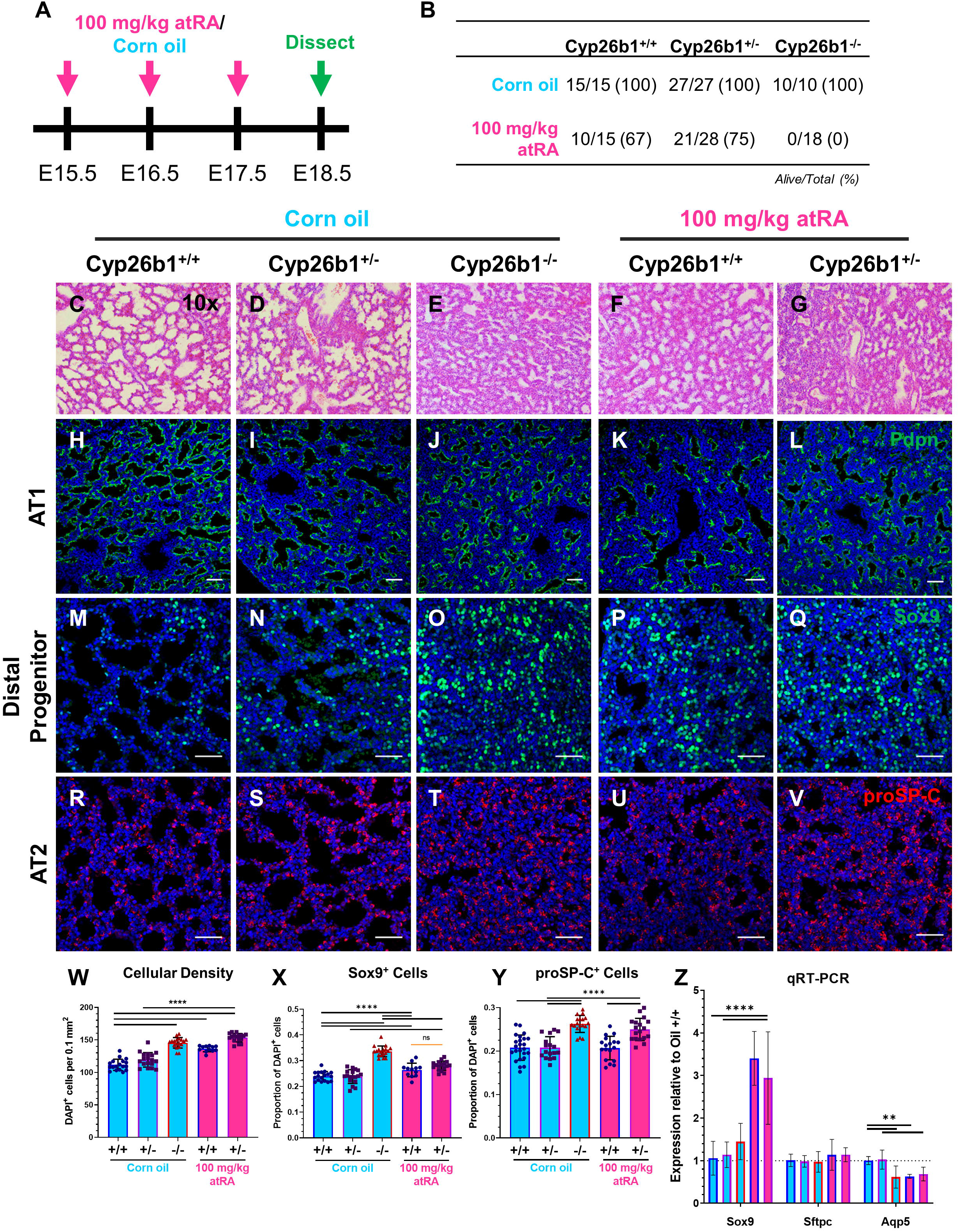
Exogenous atRA partially phenocopies loss of Cyp26b1. A) Experimental design for experiments with exogenous atRA. 100 mg/kg atRA (magenta) or equivalent dose of corn oil (control, cyan) will be gavaged to pregnant dams at E15.5, E16.5, and E17.5 and dissected at E18.5. B) Survival chart for embryos at the time of dissection for each genotype and treatment option. C-G) H&E stains of control (C-E), and atRA-treated (F-G) lungs at 10x magnification. H-L) IF stain for the AT1 cell marker Pdpn in control (H-J) and atRA-treated (K-L) lungs. M-Q) IF stain for the distal epithelial progenitor marker Sox9 in control (M-O) and atRA-treated (P-Q) lungs. IF stain for AT2 cell marker proSP-C in control (R-S) and atRA-treated (U-V) lungs. W) Number of DAPI^+^ cells per 0.1 mm^2^ demonstrating increased cellularity in the control Cyp26b1^−/−^ and atRA-treated Cyp26b1^+/+^ and Cyp26b1^+/−^ lungs. Outline color corresponds to genotype (blue = Cyp26b1^+/+^, purple = Cyp26b^+/−^, red = Cyp26b1^−/−^) and fill color corresponds to treatment group (cyan = control, magenta = atRA) X) Quantification of Sox9^+^ cells in M-Q following the same coloring scheme in W. Y) Quantification of proSP-C^+^ cells in R-V following the same coloring scheme in W. Z) qRT-PCR for Sox9, Sftpc, and Aqp5 in control and atRA-treated lungs following the same coloring scheme in W. Scale bar = 50 μm. Significance in W-Y was determined using one-way ANOVA with Tukey multiple comparison test. Significance in Z was determined using two-way ANOVA with Sidak multiple comparison test. \*\**P*<0.01, \*\*\*\**P*<0.0001.

### Exogeneous RA and loss of Cyp26b1 exhibit distinct transcriptional responses

To further validate these data, we performed qRT-PCR to see whether RA treatment induces the same changes in transcription as observed in Cyp26b1^−/−^ lungs. Consistent with previous results, exogenous atRA treatment induced expression of RARβ, Stra6, and Cyp26b1 (**Supplemental Fig. 9F, G**). We also observed a small, but significant, increase in Cyp26a1 (**Supplemental Fig. 9G**). Next, we assayed for changes in markers of each distal epithelial population. Whereas expression levels of Sftpc, Aqp5, Pdpn, Ager, and Hopx in atRA-treated lungs mirrored that of Cyp26b1^−/−^ lungs, Sox9 expression was increased and CCSP expression was decreased in atRA-treated lungs only (**Fig. 6Z**, **Supplemental Fig. 9H**). These data raise the possibility that exogenous RA may induce additional transcriptional changes not observed in Cyp26b1^−/−^ lungs. Indeed, qRT-PCR analysis of potential signaling pathways downstream of RA reveal differential expression of several Shh, Wnt, and Tgfβ members that are not differentially expressed in control Cyp26b1^−/−^ lungs (**Fig. 7**). Specifically, we found upregulation of Gli2, Gli3, Wnt4, Tgfb2, and Tgfb3 in atRA-treated lungs compared to Cyp26b1^−/−^ lungs. Additionally, several other genes – namely, Shh, Ptch1, Wnt2, Tgfbr2, Tgfbr3, and Id1 – were not differentially expressed between atRA-treated and Cyp26b1^−/−^ lungs but showed significant differences when the atRA-treated lungs were compared to their genotyped-matched controls. These data show that exogenous administration of atRA and loss of Cyp26b1 both lead to similar morphogenetic effects on lung development but do so through separate transcriptional mechanisms.

**Figure 7.**
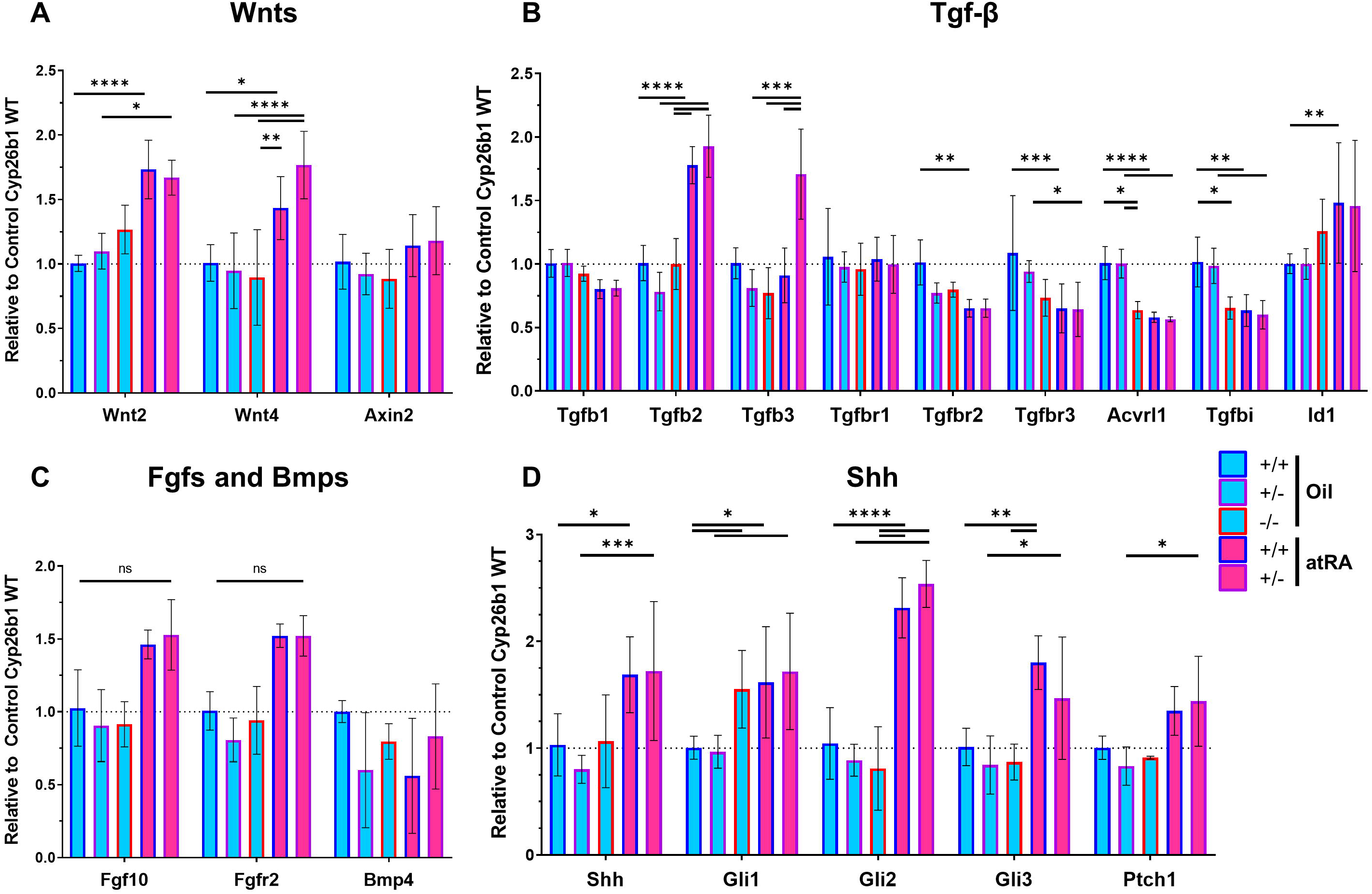
Exogenous atRA induces distinct transcriptional changes compared to Cyp26b1^−/−^ lungs. A) qRT-PCR for the Wnt pathway genes Wnt2, Wnt4, and Axin2. Outline color corresponds to genotype (blue = Cyp26b1^+/+^, purple = Cyp26b^+/−^, red = Cyp26b1^−/−^) and fill color corresponds to treatment group (cyan = control, magenta = atRA). B) qRT-PCR for members of the TGF-β signaling pathway following the same coloring scheme in A. C) qRT-PCR for established regulators of lung branching following the same coloring scheme in A. D) qRT-PCR for members of the Shh pathway following the flooring scheme in A. Significance was determined using two-way ANOVA with Sidak multiple comparison test. \**P*<0.05, \*\**P*<0.01, \*\*\**P*<0.001, \*\*\*\**P*<0.0001.

## DISCUSSION

Here we identify a novel role for Cyp26b1 in lung development. Cyp26b1 is enriched in lung ECs throughout development and loss of Cyp26b1 leads to profound changes in distal epithelial differentiation and morphogenesis, resulting in neonatal lethality. Cyp26b1^−/−^ lungs exhibit an expansion of early distal progenitors marked by both Sox9 and Lamp3/proSP-C at the expense of mature gas-exchanging AT1 cells. Currently, the only known function of Cyp26b1 is as a negative regulator of RA; however, our data indicate RA signaling is only partially increased in Cyp26b1^−/−^ lungs. Likewise, exogenous atRA during late lung formation does not fully phenocopy loss of Cyp26b1. Instead, we make the unexpected observation that Cyp26b1^−/−^ and atRA-treated lungs undergo overlapping, but separate, cellular and transcriptional responses. Therefore, our study both identifies a critical role for Cyp26b1-expressing ECs during lung development and suggests that Cyp26b1 has important roles in lung development beyond RA regulation.

Cyp26b1 is highly enriched in lung ECs throughout development, implicating ECs in directing the Cyp26b1^−/−^ phenotype. ECs have been shown to regulate lung development during branching morphogenesis and alveolarization (Del Moral et al., 2006; Jakkula et al., 2000; Lazarus et al., 2011). Additionally, ECs are critical in directing distal epithelial development in models of adult lung regeneration (Ding et al., 2011; Lee et al., 2014; Rafii et al., 2015). Our data suggest a new role for ECs during normal lung development in which ECs express Cyp26b1 to decrease RA signaling and promote distal epithelial differentiation. Although EC organization and specification appear unaltered in the mutants, we observed strong self-upregulation of Cyp26b1 specifically in ECs of Cyp26b1^−/−^ lungs. Interestingly, there was no change in Cyp26a1 or Cyp26c1 expression. Both Cyp26b1 and Cyp26a1 are known targets of RA (Rhinn and Dolle, 2012; Ross and Zolfaghari, 2011; Wu and Ross, 2010) but it is unclear why loss of Cyp26b1 or exogenous atRA both strongly upregulate only Cyp26b1 and not Cyp26a1. Based on these expression patterns, there may be unidentified EC-specific mechanisms that lead to upregulation of Cyp26b1 over the other Cyp26 enzymes. It must be stated that we cannot rule out that ECs exclusively drive the defects seen in Cyp26b1^−/−^ lungs as other non-EC populations that express Cyp26b1 at much lower levels may also be contributing to this phenotype.

Of note, we saw expansion of distal progenitors marked by both Sox9 and Lamp3, whereas the proportion of mature Lamp3^+^/Sox9^−^ AT2 cells was unaffected. Although death in Cyp26b1 mutants was previously characterized as “respiratory distress” (Yashiro et al., 2004), RDS is characterized by decreased surfactant production from AT2 cells. In that regard, our work stands in contrast to other studies that have also identified similar defects in distal epithelial differentiation but found a decrease in surfactant production or proSP-C^+^ AT2 cells (Compernolle et al., 2002; Hogmalm et al., 2014; Kersbergen et al., 2018; Rockich et al., 2013; Woik et al., 2014; Yang et al., 2002). Whereas the defects seen in these studies most likely arise from an inability of distal progenitor cells to first differentiate into AT2 cells, loss of Cyp26b1 appears to reduce the differentiation of distal progenitor into AT1.

RA has been previously shown to block distal epithelial differentiation in other contexts. Human pluripotent stem cell derived lung bud tip organoids containing Sox9^+^ distal progenitors can be maintained in a progenitor state when cultured with Fgf7, CHIR-99021, and RA (Miller et al., 2018). Removal of CHIR-99021 and RA lead to differentiation of these progenitors into the mature airway lineages. Although they did not test the effect of removing RA individually, this study is consistent with the model that RA inhibits differentiation of the distal airways. Similarly, hyperactive RA signaling through constitutively active RARα or RARβ lead to defects in distal airway formation (Wongtrakool et al., 2003). Interestingly, lungs with the constitutively active RARα transgene had a complete loss of AT2 and AT1 cells whereas lungs with the constitutively active RARβ transgene did contain some AT2 and AT1 cells, more closely mirroring Cyp26b1^−/−^ lungs. These data raise the question of whether the effects observed in Cyp26b1^−/−^ lungs are primarily mediated through RARβ or other RARs and nuclear receptors. Whereas these studies utilized *in vitro* systems or specifically engineered transgenic lines, we identify Cyp26b1 as an endogenous physiologic mechanism that decreases RA signaling during normal lung development.

Exogenous administration of atRA beginning at E15.5 is able to partially phenocopy loss of Cyp26b1. Although morphologic changes between atRA-treated and Cyp26b1^−/−^ lungs were indistinguishable from one another, cell fate changes in atRA-treated lungs did not fully mirror that in Cyp26b1^−/−^ lungs. One possible explanation for this is that distal epithelial progenitors are specified as early as E13.5 and may not be receptive to changes in fate by the first dose of RA at E15.5 (Frank et al., 2019). Cyp26b1^−/−^ lungs are exposed to higher levels of RA throughout lung development potentially leading to a more severe difference in lineage specification. Despite these results that suggest that RA drives the defects in lung formation, our transcriptional analyses suggest that this phenotype is modulated by an additional RA-independent mechanism. First, we saw little change in the expression of multiple signaling pathways known to be regulated by RA during lung development. Second, several genes in these pathways were differentially expressed in atRA-treated lungs. These data indicate that loss of Cyp26b1 and exogenous atRA can both result in similar gross defects in lung development, but do so through different signaling pathways. Additional studies need to be performed in order to identify these RA-independent mechanisms that drive the defects in Cyp26b1^−/−^ lungs.

From our studies, it is clear that Cyp26b1 is essential for the development of a functional lung via proper differentiation of lung progenitor cells. It is of particular interest that Cyp26b1 in the lung is primarily expressed in endothelial cells. We propose that our study identified Cyp26b1 as a novel endothelial rheostat for RA activity in embryonic organs. Given that Cyp26b1 null embryos display increased cellular density and contain an expansion of progenitors at the expense of mature gas-exchanging AT1 cells, it will be of great interest to see whether its loss causes defects in other organs. Together, our data provide new perspectives on the complex mechanisms by which RA regulates lung development and suggests novel mechanisms downstream of Cyp26b1. These studies will likely prove useful to developing therapies for newborns with lung maturation defects.

## SUPPLEMENTAL FIGURE LEGENDS

**Supplemental Figure 1. Cyp26b1 is highly enriched in lung, kidney, and other endothelial cell beds in post-natal mice.** A-C) Violin plots of Cyp26b1 expression in scRNA-seq of adult kidneys (A), lungs (B), and in all adult organs combined (C) obtained from Tabula Muris consortium (Tabula Muris et al., 2018). EC populations are highlighted in red. Data accessed through https://tabula-muris.ds.czbiohub.org/ D) Scatter plot of Drop-seq analysis of P1 lungs from LungGENS (Du et al., 2015; Du et al., 2017; Guo et al., 2019). Vascular-ECs are marked with Red arrow. Data accessed through https://research.cchmc.org/pbge/lunggens/SCLAB.html. E) Bulk RNA-seq of Tie2-GFP^+^ P7 ECs from brain, liver, lung, and kidney compared to GFP^−^ cells (Sabbagh et al., 2018). Data accessed through https://markfsabbagh.shinyapps.io/vectrdb/. F-F’) tSNE plot of scRNA-seq data of human fetal kidneys at week 16 of gestation (Hochane et al., 2019). Red arrow highlights endothelial cluster. Data accessed through https://home.physics.leidenuniv.nl/~semrau/humanfetalkidneyatlas/.

**Supplemental Figure 2. Cyp26b1 is expressed in multiple organs at E12.5.** A-E) In situ hybridization for Cyp26b1 in the head (A), hindbrain (B), forelimb (C), somites (D), and heart (E). Mouth in (A) is marked for orientation. F) Zoomed in image of endocardial cushion shown in E. Magnifications for each image are shown.

**Supplemental Figure 3. Gross histology of Cyp26b1^−/−^ lungs in late gestation**. A-F) H&E stains of Cyp26b1^+/+^ (A, C, E) and Cyp26b1^−/−^ (B, D, F) lungs at E15.5 (A-B), E16.5 (C-D), and E17.5 (E-F). Magnification for all images shown.

**Supplemental Figure 4. Validation of defects in distal epithelial differentiation using independent AT1 and AT2 cell markers**. A-C) IF stain for the AT2 markers Lamp3 and proSP-C. C) Quantification of all proSP-C^+^ cells and all Lamp3^+^ cells with respect to DAPI. D) Stratification of data shown in C into proSP-C/Lamp3 Double positive cells (first set), proSP-C single positive cells (second set), and Lamp3 single positive cells (third set) as a proportion of DAPI^+^ cells. E) Quantification of proSP-C/Lamp3 double positive cells (green) and proSP-C single positive cells (orange) with respect to all <u>proSP-C^+^</u> cells. F) Quantification of proSP-C/Lamp3 double positive cells (green) and Lamp3 single positive cells (orange) with respect to all <u>Lamp3^+^</u> cells. The proportions of proSP-C/Lamp3 double positive, proSP-C single positive, and Lamp3 single positive AT2 cells are the same between Cyp26b1^+/+^ and Cyp26b1^−/−^ lungs. Greater than 90% of all AT2 cells are proSP-C^+^/Lamp3^+^. G, H) IF stain for the AT1 marker Pdpn. I) qRT-PCR analysis for AT1 markers Pdpn, HOPX, and Ager and the AT2 marker Lamp3. J) Quantification of Lamp3/Sox9 double positive cells (green) and Sox9 single positive cells (orange) with respect to all <u>Sox9^+^</u> cells. K) Quantification of Lamp3/Sox9 double positive cells (green) and Lamp3 single positive cells (orange) with respect to all <u>Lamp3^+^</u> cells. Scale bar = 50 μm. Significance was determined using two-way ANOVA with Sidak multiple comparison test. \**P*<0.05, \*\*\*\**P*<0.0001.

**Supplemental Figure 5. Cyp26b1^−/Δ10^ lungs phenocopy Cyp26b1^−/−^ lung defects in increased cellular density and distal epithelial differentiation.** A-B) E18.5 Cyp26b1^+/+^ and Cyp26b1^−/Δ10^ embryos at dissection. C-D) H&E stain of E18.5 Cyp26b1^+/+^ and Cyp26b1^−/Δ10^ lungs at 10x. E-G) IF stain for the Sox9^+^ distal progenitors and quantification relative to DAPI^+^ cells. H-J) IF stain for the proSP-C^+^ AT2 cells and quantification relative to DAPI^+^ cells. K-L) IF stain for the Pdpn^+^ AT1 cells demonstrating smaller distal airspaces. M) Number of DAPI^+^ cells per 0.1 mm^2^ demonstrating increased cellularity in the transheterozygotes. Scale bar = 50 μm. Significance was determined using Student’s T-Test. \*\*\*\**P*<0.0001.

**Supplemental Figure 6. Proximal airways are unaffected in Cyp26b1^−/−^ lungs.** A-B) IF stains for CCSP (red) and FoxJ1 (white) in E18.5 Cyp26b1^+/+^ (A) and Cyp26b1^−/−^ (B) lungs. C) Flow charts for analysis of BASC populations. D) Quantification of the frequency BASCs (CD31^−^/CD45^−^/Sca1^+^/Ep-CAM^+^) as a proportion of all live cells in E16.5 – E18.5 lungs. Quantifications are standardized to the frequency of BASCs in Cyp26b1^+/+^ samples per experiment. Scale bar = 50 μm.

**Supplemental Figure 7. Stromal and Vascular lineages are unaffected in Cyp26b1^−/−^ lungs.** A-B) If stain for the stromal markers Pdgfr-α (green) and Pdgfr-β (red) in E18.5 lungs. C-D) IF stain for the broad EC marker VE-Cad (green). E-F) IF stain for VE-Cad (green), Pdgfr-β (red), and E-Cad (white) to examine pericyte-EC interaction. G-H) IF stain for Emcn (green), Claudin-5 (red), and Sox17 (white) to differentiate arterial vs venous differentiation. A = artery, V = vein, Br = bronchi/bronchiole. Emcn is expressed in all but arterial ECs while Claudin-5 and Sox17 are more highly expressed in arterial ECs compared to capillary and venous ECs. I-J) If stain for the lymphatic marker Lyve1 (red) and Emcn (white). K-L) IF stain for VE-Cad (green), Sm22a (magenta), and E-Cad (white). Scale bar = 50 μm (A-D,G-L), 25 μm (E-F).

**Supplemental Figure 8. Other signaling pathways implicated in lung development are unaffected in Cyp26b1^−/−^ lungs.** A) qRT-PCR for members of the Tgf-β signaling pathway. B) qRT-PCR for established regulators of lung branching: Fgf10, Fgfr2, and Bmp4. C) qRT-PCR for members of the Wnt signaling pathway. D) qRT-PCR for members of the Shh signaling pathway. Expression levels are standardized to that in Cyp26b1^+/+^ lungs for each gene. Significance was determined using two-way ANOVA with Sidak multiple comparison test. \*\*\*\**P*<0.0001.

**Supplemental Fig. 9. Morphologic and transcriptional changes in RA and epithelial genes with atRA treatment**. A-E) Lungs from E18.5 control (A-C) and atRA-treated (D-E) Cyp26b1^+/+^, Cyp26b1^+/−^, and Cyp26b1^−/−^ lungs at time of dissection. Magnified views of the left lobe are shown in A’, B’, C’, D’, and E’ to highlight loss of distal airspaces in Cyp26b1^−/−^ and atRA-treated Cyp26b1^+/+^ and Cyp26b1^+/−^ lungs. F) qRT-PCR for RARβ and Stra6. Outline color corresponds to genotype (blue = Cyp26b1^+/+^, purple = Cyp26b^+/−^, red = Cyp26b1^−/−^) and fill color corresponds to treatment group (cyan = control, magenta = atRA). G) qRT-PCR for Cyp26b1 and Cyp26a1 following the same coloring scheme in F. H) qRT-PCR for the epithelial genes Pdpn, Ager, HOPX, and CCSP following the same coloring scheme in F. Scale bar = 1 mm. Significance was determined using two-way ANOVA with Sidak multiple comparison test. \**P*<0.05, \*\**P*<0.01, \*\*\**P*<0.001, \*\*\*\**P*<0.0001.

